# Identification of novel hepaciviruses in rock pigeon (*Columba livia* [Gmelin, 1789]), rusty-margined flycatcher (*Myiozetetes cayanensis* [Linnaeus, 1766]), and Hispaniolan amazon (*Amazona ventralis* [Statius Muller, 1776])

**DOI:** 10.64898/2026.03.05.709806

**Authors:** Sae Kawano, Mai Kishimoto, Sakiho Imai, Tomohisa Tanaka, Kohji Moriishi, Masayuki Horie

## Abstract

Recent advances in sequencing technology and transcriptome mining have revealed highly divergent hepaciviruses in birds. However, only a limited number of avian hepaciviruses have been identified to date, leaving their diversity and evolutionary history poorly understood. Moreover, deep phylogenetic gaps among known avian hepaciviruses suggest that additional lineages remain undiscovered. Here, we screened publicly available RNA-seq data and identified three previously undescribed hepaciviruses from rock pigeon (*Columba livia*), rusty-margined flycatcher (*Myiozetetes cayanensis*), and Hispaniolan amazon (*Amazona ventralis*), named rock pigeon hepacivirus (RpHV), rusty-margined flycatcher hepacivirus (RfHV), and Hispaniolan amazon hepacivirus (HaHV). Although these three viruses meet the ICTV species demarcation criteria relative to their closest known relatives, the NS5B-based criterion was not satisfied between RfHV and HaHV. Notably, however, their genome sequence identity is low at 43.2%, and their hosts differ at the order level, suggesting that their classification warrants further consideration. Our phylogenetic analysis showed that avian hepaciviruses, including those found in this study, are monophyletic, but phylogenetic incongruence was observed between avian hepaciviruses and their hosts, suggesting past cross-species transmission among avian hepaciviruses. Overall, this study provides novel insights into the diversity and evolution of hepaciviruses.

## Introduction

Hepaciviruses (genus *Hepacivirus*, family *Flaviviridae*) are positive-sense single-stranded RNA viruses [1]. Their genomes are approximately 8.9–10.5 kb in length and encode a single polyprotein of ∼3,000 amino acids, flanked by untranslated regions (UTRs) at both ends. In hepatitis C virus (HCV), a representative virus in the genus *Hepacivirus*, the polyprotein is processed into structural proteins (core, E1, and E2), p7, and nonstructural proteins NS2–NS5B. After the establishment of the genus *Hepacivirus* in 1996, HCV had been the only recognized species for many years in the genus. GB virus B (GBV-B), which was identified in New World monkeys in 1995 [2], was initially not classified within the genus because of substantial sequence divergence. Since 2011, high-throughput sequencing has markedly expanded the known diversity of hepaciviruses in mammals, with viruses identified in multiple host taxa including dogs [3], horses [4], bats [5], rodents [6–8], non-human primates [9], and cattle [10, 11]. Consequently, the taxonomy of the genus has been updated as additional lineages have been characterized [12]. Additionally, more divergent hepaciviruses have been reported in a broader range of vertebrates, including fish [13], reptiles [14], and birds [15–21], although they have not yet been officially classified into the genus *Hepacivirus* [1].

Avian hepaciviruses were first described in 2019 during an etiological investigation of severe disease in ducks (*Anas platyrhynchos*) [15], with subsequent identification in ducks [18], bald eagles (*Haliaeetus leucocephalus*) [16], and Ōkārito Rowi (*Apteryx rowi*) [21]. Additionally, hepacivirus-like sequences have been reported in the common banded mosquito (*Culex annulirostris*), suspected to originate from tawny frogmouths (*Podargus strigoides*) [17]. Furthermore, partial hepacivirus sequences were also detected from zebra finches (*Taeniopygia guttata*) [19] and Fiordland penguin (tawaki) (*Eudyptes pachyrhynchus*) [20]. Although these viruses have occasionally been detected in symptomatic hosts [15, 16], it is unclear whether the hepaciviruses are associated with these diseases. Notably, large phylogenetic gaps among known avian hepaciviruses exist, strongly suggesting the presence of undiscovered hepaciviruses [21].

Therefore, further investigation is necessary to understand the diversity and evolution of avian hepaciviruses and to characterize them.

In this study, we screened public RNA-seq data from birds to identify novel avian hepaciviruses. We identified three novel hepaciviruses, named rock pigeon hepacivirus (RpHV) from *Columba livia* (Gmelin, 1789), rusty-margined flycatcher hepacivirus (RfHV) from *Myiozetetes cayanensis* (Linnaeus, 1766), and Hispaniolan amazon hepacivirus (HaHV) from *Amazona ventralis* (Statius Muller, 1776) and characterized these viruses.

## Materials and Methods

### Screening of public RNA-seq data derived from birds

A total of 42,122 avian RNA-seq datasets in the NCBI Sequence Read Archive (SRA) [22] (Table S1) were screened for sequence similarity against known avian hepaciviruses using SRAminer pre0.1 [23] with the option “-N 1000000”. For RNA-seq datasets where at least one read was hit by SRAminer, the top-hit read was extracted. Using the extracted reads as queries, a sequence similarity search was performed against the NCBI clustered nr database (in April 2024) [22] using BLASTx 2.15.0+ [24] with the options “-evalue 0.05 -max_target_seqs 10 -word_size 2 -seg no -lcase_masking”. RNA-seq datasets with reads whose top BLASTx hits were avian hepaciviruses were selected for further analysis.

### Identification of avian hepacivirus-like contigs

The selected RNA-seq datasets were downloaded and preprocessed using fastp 0.23.2 [25] with the options “-x -y -l 35”. The genome sequence of the host species corresponding to each RNA-seq dataset, or a congeneric species if the host genome was unavailable, was downloaded from Genomesync [26]. Subsequently, the preprocessed reads were mapped to the corresponding host genomes using HISAT2 version 2.2.1 [27], and unmapped reads were extracted using SAMtools 1.16.1 [28].

The extracted reads were *de novo* assembled using Trinity v2.15.1 [29] with the default settings. The resulting contigs were subjected to a two-step sequence similarity search using BLASTx 2.15.0+ as follows. The first step involved searching against a custom database consisting of known avian hepacivirus sequences. This custom database was created using manually curated avian hepacivirus sequences (Table S2). The search options were “-evalue 1e-10 -max_target_seqs 10 -word_size 2 -seg no -lcase_masking”. For contigs yielding hits in the first step, a second search was performed against the NCBI clustered nr database (in June 2024). The options were “-evalue 0.5 -max_target_seqs 10 -word_size 2 -seg no -lcase_masking”. Contigs for which the top hits corresponded to avian hepaciviruses were identified as avian hepacivirus-like contigs. In cases where multiple partial avian hepacivirus-like contigs were obtained from the same RNA-seq dataset, they were assembled using the Geneious Assembler in Geneious Prime Version 2025.0.3 (https://www.geneious.com/) with the default settings. Avian hepacivirus-like contigs (Table S3) of 5,000 nucleotides or longer were subjected to subsequent analyses.

### Contig validation and ORF prediction

Original RNA-seq reads were mapped back to the corresponding hepacivirus-like contigs using Magic-BLAST 1.6.0 [30]. Read depth at each position was manually inspected using Geneious Prime, and terminal regions with a read depth of 1 were trimmed. Within the putative coding regions, positions with a read depth of 2 or more, or those with a read depth of 1 with a sequencing quality score of 37 or higher, were considered reliable, and “unreliable positions” were trimmed. The trimmed contigs were used for subsequent analyses as novel avian hepaciviruses.

Based on the ORF lengths of known avian hepaciviruses, ORFs of 5,000 nucleotides or longer were searched for within the contigs using Geneious Prime.

### Species demarcation analysis

Each avian hepacivirus sequence was pairwise aligned with the amino acid sequence of HCV genotype 1a H77 (AF011751.1) using MAFFT 7.490 with the E-INS-i algorithm [31]. Then, conserved regions within NS3 (corresponding to amino acid positions 1,123–1,566 of HCV genotype 1a H77) and NS5B (corresponding to amino acid positions 2,536–2,959 of HCV genotype 1a H77), as defined by the ICTV, were extracted. The extracted sequences were then aligned using MAFFT, and then pairwise sequence identities were determined using Sequence Demarcation Tool version 1.2 [32]. Each *p*-distance was calculated based on the identity.

### Screening of novel hepaciviruses in public RNA-seq datasets

First, the avian hepacivirus-like contigs obtained above (Table S3) were re-analyzed. Contigs with a length of 1,000 nucleotides or longer were aligned with the nucleotide sequences of the novel hepaciviruses using MAFFT with the E-INS-i algorithm to extract the conserved NS3 and NS5B regions as described above. The extracted regions were translated, and pairwise alignment was performed with the corresponding novel hepaciviruses to calculate the *p*-distance. Contigs exhibiting an NS3 amino acid p-distance of 0.25 or less, or an NS5B amino acid *p*-distance of 0.3 or less relative to the corresponding novel hepaciviruses, were classified as conspecific with the corresponding novel hepaciviruses, which were further analyzed as follows. The contig sequences were assessed by back-mapping of original reads as described above.

Pairwise alignment was performed using MAFFT with the E-INS-i algorithm to calculate the nucleotide sequence identity. For partial sequences, multiple alignment including a full-length sequence was performed using MAFFT with the E-INS-i algorithm, and the average nucleotide sequence identities within the aligned regions was calculated.

Next, a total of 42,122 avian public RNA-seq datasets (Table S1) were screened for sequence similarity against the sequences of the discovered novel hepaciviruses using SRAminer pre0.1 and Magic-BLAST 1.6.0. The option “-N 1000000” was used for SRAminer, while the default settings were employed for Magic-BLAST. For RNA-seq datasets where at least one read was hit, the top-hit reads from SRAminer, and Magic-BLAST were used as queries for a sequence similarity search against the NCBI clustered nr database (in January 2025) using BLASTx 2.15.0+. The options were set to “-evalue 0.05 -max_target_seqs 10 -word_size 2 -seg no -lcase_masking”. Datasets in which the top hit corresponded to an avian hepacivirus were identified as containing novel hepacivirus-like reads.

### Gene annotation of novel hepaciviruses

RpHV and RfHV were pairwise aligned with their closest relatives using MAFFT with the E-INS-i algorithm. Then, the RpHV and RfHV genomes were annotated based on the annotations of their closest relatives. For HaHV, pairwise alignment with RfHV was performed using QIAGEN CLC Genomics Workbench 25.0.2 (https://digitalinsights.qiagen.com/), and annotations were assigned based on RfHV.

### Phylogenetic analyses

The putative amino acid sequences of the three novel hepaciviruses and 33 representative hepaciviruses were aligned using MAFFT with the E-INS-i algorithm. Ambiguously aligned regions were trimmed using trimAl 1.4.rev22 [33] with the “-strict” option. A phylogenetic tree was inferred by the maximum likelihood method using RAxML Next Generation 1.1.0 [34] with the LG+I+G4+FC model selected using ModelTest-NG 0.2.0 [35]. The reliability of the tree was evaluated by the transfer bootstrap method [36] with 1,000 replicates.

Phylogenetic trees were also inferred using the conserved NS3 or NS5B regions of avian hepaciviruses. Each known avian hepacivirus sequence was pairwise aligned with the amino acid sequence of HCV genotype 1a H77 using MAFFT with the E-INS-i algorithm to extract the NS3 and NS5B regions. The extracted sequences were multiple aligned, which were used for phylogenetic inference. Phylogenetic trees were reconstructed by the maximum likelihood method with the LG+I+G4+F and LG+G4 models for the NS3 and NS5B regions, respectively. The reliabilities of trees were evaluated as described above.

To compare the phylogenetic tree of avian hepaciviruses with that of their hosts, a phylogenetic tree of the host species was retrieved from TimeTree 5 [37].

All trees were visualized and edited using MEGA12 [38].

### Sequence data of novel hepaciviruses

All sequences of the novel hepaciviruses identified in this study were deposited in the DNA Data Bank of Japan (DDBJ) under the accession numbers BR002552–BR002563. Since the RNA-seq datasets SRR13708282 and SRR22541704, and SRR13685388 and SRR22541710, respectively contained identical sequence data, only the contigs obtained from SRR13708282 and SRR13685388 were deposited (Table S4).

## Results

### Identification of three novel hepaciviruses from public RNA-seq data

To identify novel avian hepaciviruses, we comprehensively analyzed public avian RNA-seq data. Consequently, we detected many hepacivirus-like contigs (Table S3), including the below three novel hepaciviruses (Fig. 1A and Table 1). We identified the first virus from RNA-seq data derived from the intraperitoneal fat of a rock pigeon *(Columba livia*) (SRR21669561), named rock pigeon hepacivirus (RpHV). The contig (BR002552) is 9,905 nucleotides (nt) in length and contains a single open reading frame (ORF) putatively encoding a polyprotein of 3,045 amino acids (Fig. 1A). This contig exhibited 53.3% nucleotide sequence identity with its closest relative, Jogalong virus (MN133813.1). We also identified near-full-length RpHV sequences from two other RNA-seq datasets (BR002562 and BR002563), but we selected the longest contig (BR002552) as the representative sequence for subsequent analyses. Regarding ORF annotation, although we observed an upstream start codon in the representative contig (BR002552) relative to the ORFs of the other two contigs (BR002562 and BR002563), we adopted the start codon shared among all three contigs. Additionally, through re-analysis of contigs and screening for RpHV in public RNA-seq data, we detected seven partial RpHV sequences from an additional three datasets (Fig. 1B). Please note that the reads in SRR13708282 and SRR13685388 datasets are identical to those in SRR22541710 and SRR22541704, respectively, and therefore we only analyzed each former dataset. The average nucleotide sequence identity among the RpHVs ranged from 83.5% to 99.9% (Fig. 1C).

**Figure 1.**
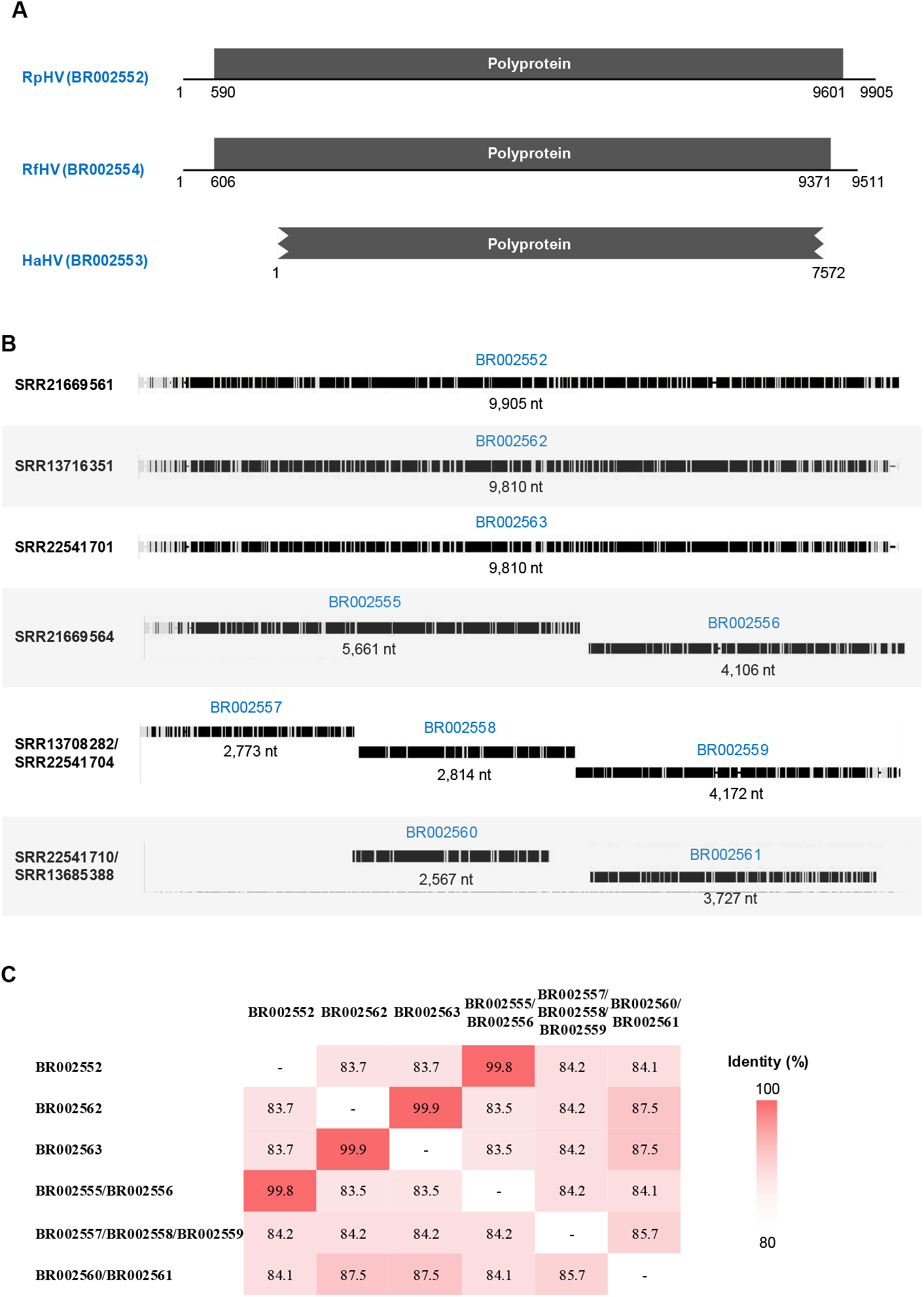
Genomes of novel hepaciviruses. **(A)** Schematic diagrams of genome organization of novel hepaciviruses. RpHV, rock pigeon hepacivirus; RfHV, rusty-margined flycatcher hepacivirus; HaHV, Hispaniolan amazon hepacivirus. Numbers indicate nucleotide positions. Black boxes represent ORFs. Partial ORF is indicated by jagged lines. **(B)** Schematic representation of contig sequences of rock pigeon hepacivirus (RpHV). Multiple sequence alignment of all contigs is shown. Black-shaded areas indicate regions where nucleotides match the consensus sequence, while white areas indicate regions where nucleotide does not match. The accession number of the source RNA-seq dataset is shown to the left of each contig sequence. The DDBJ accession numbers are shown above, and the contig lengths are shown below the boxes. Heatmap showing nucleotide sequence identities (%) among RpHVs.

**Table 1.**
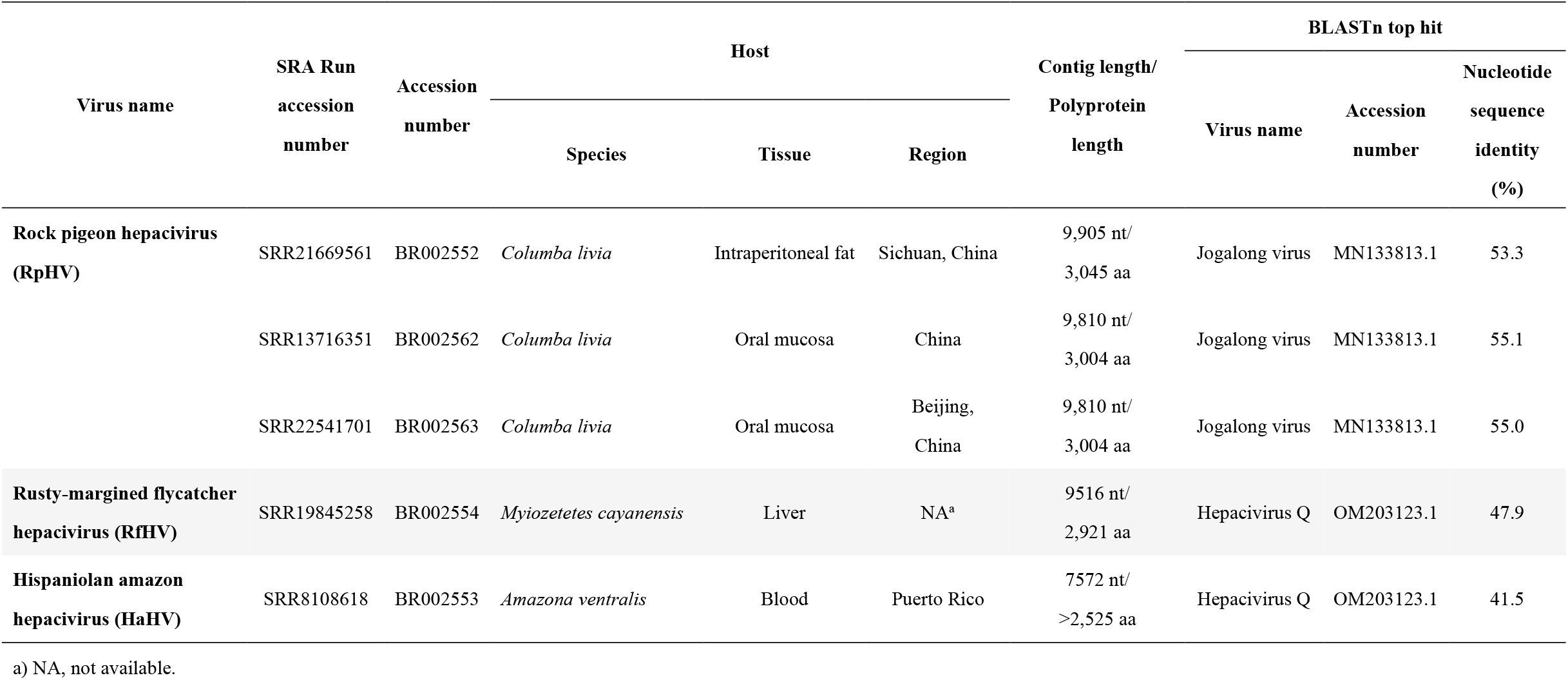
Summary of representative hepaciviruses identified in this study.

| Virus name | SRA Run<br>accession<br>number | Accession<br>number | Host |  |  | Contig length/<br>Polyprotein<br>length | BLASTn top hit |  |  |
| --- | --- | --- | --- | --- | --- | --- | --- | --- | --- |
|  |  |  | Species | Tissue | Region |  | Virus name | Accession<br>number | Nucleotide<br>sequence<br>identity<br>(%) |
| Rock pigeon hepacivirus<br>(RpHV) | SRR21669561 | BR002552 | *Columba livia* | Intraperitoneal fat | Sichuan, China | 9,905 nt/<br>3,045 aa | Jogalong virus | MN133813.1 | 53.3 |
|  | SRR13716351 | BR002562 | *Columba livia* | Oral mucosa | China | 9,810 nt/<br>3,004 aa | Jogalong virus | MN133813.1 | 55.1 |
|  | SRR22541701 | BR002563 | *Columba livia* | Oral mucosa | Beijing,<br>China | 9,810 nt/<br>3,004 aa | Jogalong virus | MN133813.1 | 55.0 |
| Rusty-margined flycatcher<br>hepacivirus (RfHV) | SRR19845258 | BR002554 | *Myiozetetes cayanensis* | Liver | NA<sup>a</sup> | 9516 nt/<br>2,921 aa | Hepacivirus Q | OM203123.1 | 47.9 |
| Hispaniolan amazon<br>hepacivirus (HaHV) | SRR8108618 | BR002553 | *Amazona ventralis* | Blood | Puerto Rico | 7572 nt/<br>>2,525 aa | Hepacivirus Q | OM203123.1 | 41.5 |

a) NA, not available.

We identified the second hepacivirus in RNA-seq data derived from the liver of a rusty-margined flycatcher (*Myiozetetes cayanensis*) (SRR19845258), named rusty-margined flycatcher hepacivirus (RfHV). The contig (BR002554) is 9,516 nt in length and contains an ORF encoding a polyprotein of 2,921 amino acids (Fig. 1A). This contig exhibited 47.9% nucleotide sequence identity with its closest relative, Hepacivirus Q (OM203123.1). Although we attempted to detect further RfHV-positive data, we did not detect RfHV-derived reads in any RNA-seq datasets.

We identified the third hepacivirus in RNA-seq data derived from the blood of a Hispaniolan amazon (*Amazona ventralis*) (SRR8108618), named Hispaniolan amazon hepacivirus (HaHV). The contig (BR002553) is 7,572 nt in length, which corresponds to a partial ORF spanning from the latter half of the E2 region to the first half of the NS5B region (Fig. 1A). This contig exhibited 41.5% nucleotide sequence identity with its known closest relative, Hepacivirus Q (OM203123). Despite our efforts to detect further HaHV-positive data, we did not detect HaHV in any RNA-seq datasets.

### Novel hepaciviruses could represent new species within the genus Hepacivirus

We evaluated the taxonomic status of the identified viruses by calculating amino acid *p*-distances for the NS3 and NS5B regions (Fig. 2), in accordance with the ICTV species demarcation criteria (*p*-distance of >0.25 for the NS3 and >0.3 for the NS5B regions to the closest relative) [1]. RpHV exhibited *p*-distances of 0.34 (NS3) and 0.35 (NS5B) to its closest relative, Jogalong virus.

**Figure 2.**
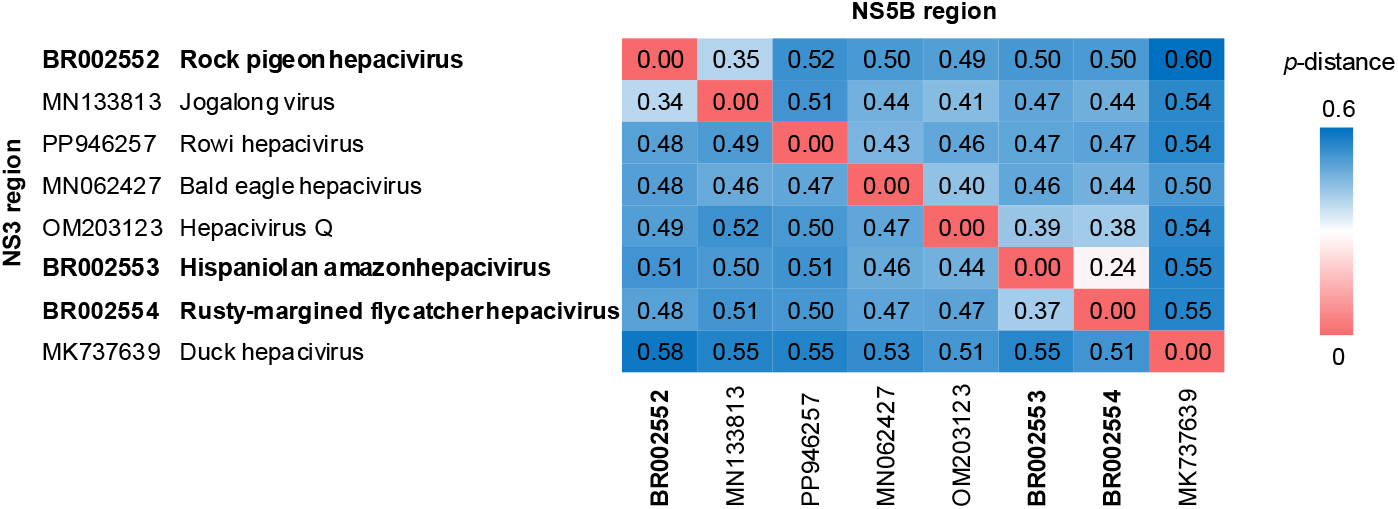
Pairwise identities of conserved NS and NS5B regions among avian hepaciviruses. Heatmap of pairwise amino acid identities in the conserved NS3 and NS5B regions of avian hepaciviruses. The lower and upper triangles show identities for the NS3 and NS5B regions, respectively.

Similarly, RfHV and HaHV showed substantial divergence from their closest relative, Hepacivirus Q, with *p-*distances of 0.47 and 0.44 in the NS3 region, and 0.37 and 0.39 in the NS5B region, respectively. These values meet the ICTV species demarcation criteria. Note that, however, the *p*-distances between RfHV and HaHV are 0.37 for NS3 and 0.24 for NS5B, latter of which does not meet the ICTV criteria (see Discussion).

### Characterization of novel hepaciviruses

To characterize the identified novel hepaciviruses, we analyzed the metadata of virus-positive datasets. This revealed that the RpHV-positive RNA-seq datasets originated from three BioProjects (Tables 1 and S5). These data are derived from intraperitoneal fat, subcutaneous fat, and oral mucosa samples of rock pigeons in China. Of these, six datasets were derived from clinically healthy individuals, while two were from individuals exhibiting signs of disease, the details of which are unknown (Tables 1 and S5). However, it should be noted that though RNA-seq datasets SRR13708282 and SRR22541704 contains identical reads as described above, their descriptions of health status are different, which compromises the reliability of the data.

The RfHV-positive RNA-seq dataset is derived from a liver sample of a rusty-margined flycatcher. The RNA-seq dataset in which HaHV was detected was derived from a blood sample of a Hispaniolan amazon kept in Puerto Rico. No information regarding the health status of the hosts was available for RfHV and HaHV (Table 1s and S5).

### Phylogenetic analyses of avian hepaciviruses

To understand the evolutionary relationship between the novel hepaciviruses and others, we performed phylogenetic analysis using the putative polyprotein sequences (Fig. 3). The tree showed that novel avian hepaciviruses are clustered with the previously known avian hepaciviruses. While RpHVs are clustered with Jogalong virus, RfHV and HaHV are clustered together with Hepacivirus Q.

**Figure 3.**
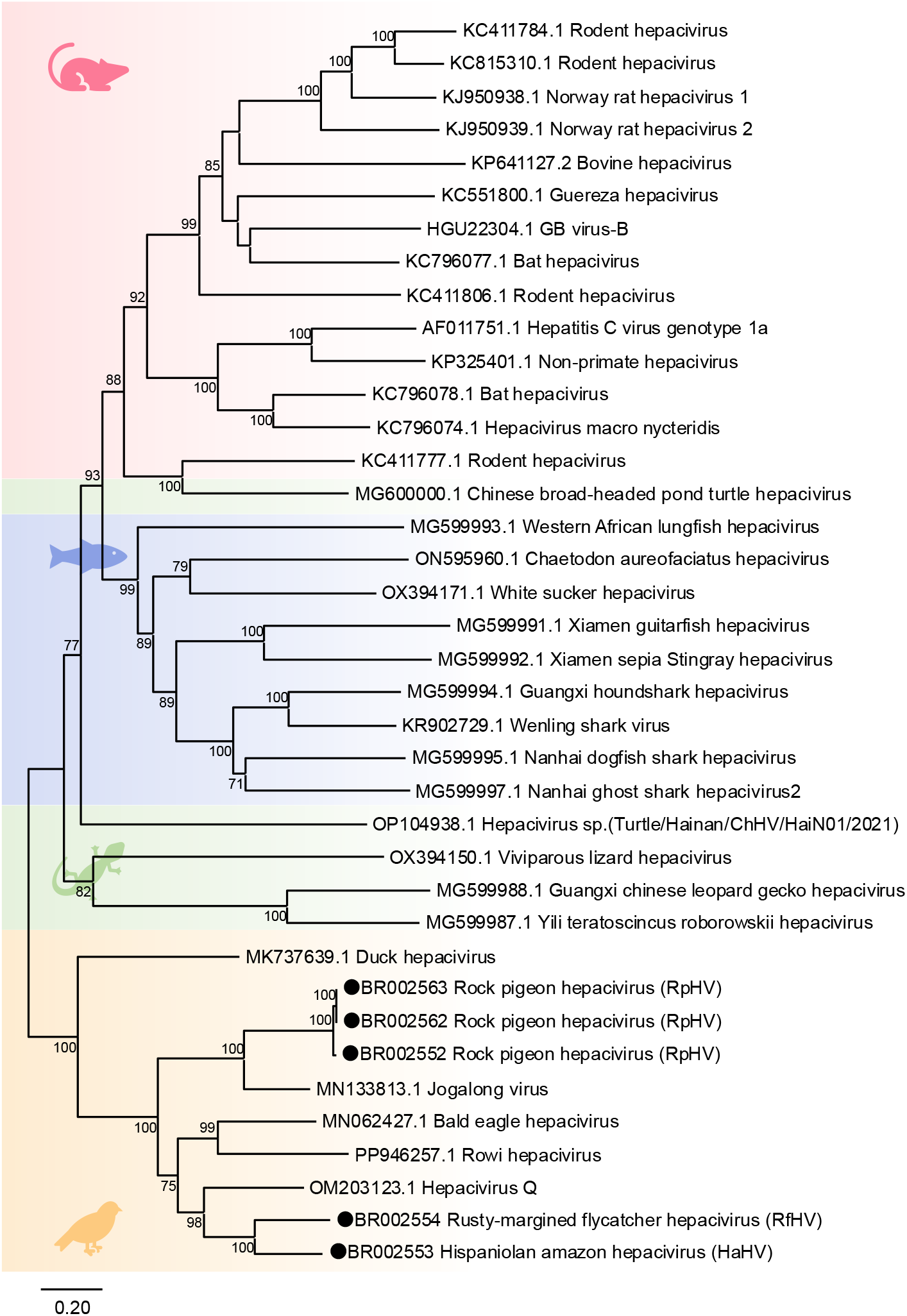
Phylogenetic tree of hepaciviruses. A maximum-likelihood phylogenetic tree was constructed using polyprotein amino acid sequences of the three novel hepaciviruses identified in this study and representative hepaciviruses. The scale bar indicates the number of amino acid substitutions per site. Black circles indicate the novel hepaciviruses identified in this study.

To further elucidate the evolutionary relationships among avian hepaciviruses, we conducted phylogenetic analyses using conserved amino acid sequences of the NS3 and NS5B regions, including avian hepaciviruses with available partial genome sequences (Fig. S1). Interestingly, the topology of the trees was different, suggesting past recombination events within avian hepaciviruses in accordance with a previous report [18]. We also performed recombination analysis using RDP4 [39], but could not detect clear recombination breakpoints, which is likely due to the high genetic divergence of the sequences used for the analysis.

### Incongluence of viral and host phylogeny

To further understand the evolution of avian hepaciviruses, we compared the topologies of trees between avian hepaciviruses and their hosts (Fig. 4). We found some incongruences between those trees, suggesting the past cross-species transmission of avian hepaciviruses (see Discussion).

**Figure 4.**
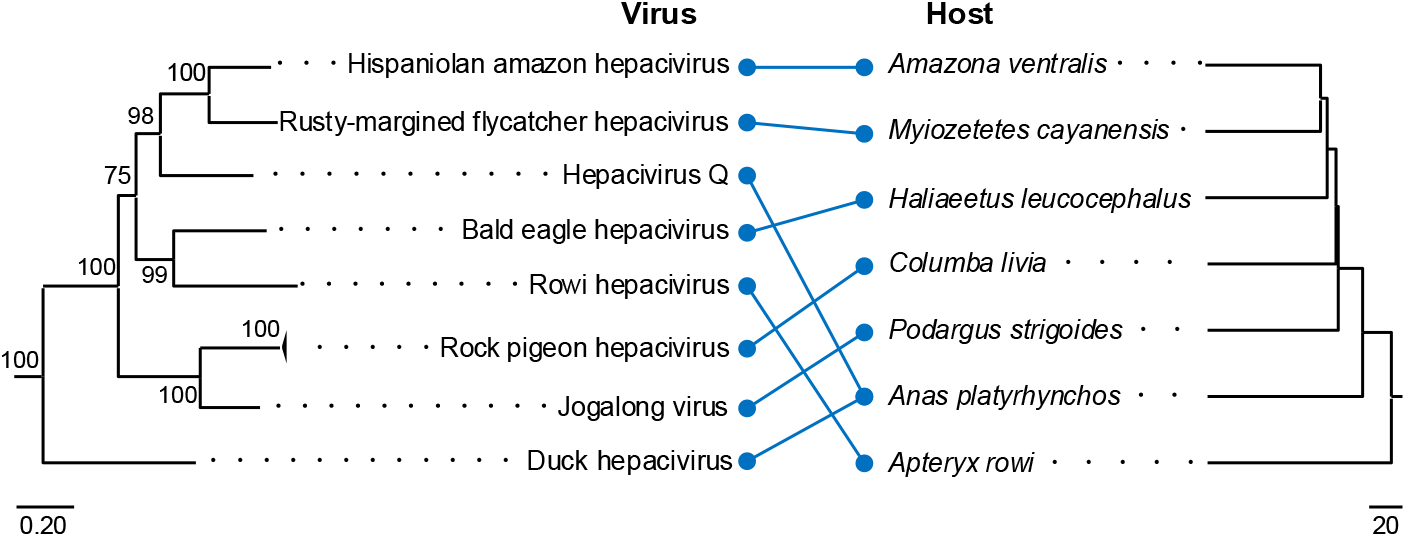
Comparison of phylogenetic trees of avian hepaciviruses and their hosts. The left panel shows the phylogenetic tree of avian hepaciviruses, and the right panel shows the phylogenetic tree of their hosts. Viruses and their corresponding hosts are connected by blue lines. The phylogenetic tree of avian hepaciviruses was adapted from Figure 3, and the host phylogenetic tree was obtained from TimeTree 5. The left scale bar indicates the number of amino acid substitutions per site. The right scale bar represents time in millions of years.

## Discussion

Although hepaciviruses have been discovered in a wide range of vertebrates, the diversity and evolutionary history of avian hepaciviruses remain poorly understood. In this study, we identified three novel hepaciviruses, designated RpHV, RfHV, and HaHV, from bird species not previously recognized as hepacivirus hosts, namely the rock pigeon, rusty-margined flycatcher, and Hispaniolan amazon. Because these viruses meet the ICTV species demarcation criteria [1], we propose them as novel species within the genus *Hepacivirus*. Thus, this study contributes to a deeper understanding of the diversity and evolution of avian hepaciviruses.

While strictly applying the current ICTV species demarcation criteria would classify RfHV and HaHV as the same species, they may be classified as two distinct species considering differences in their hosts and genetic distance. Although the amino acid *p*-distance between RfHV and HaHV in the NS3 region (0.37) meets the ICTV criterion, the *p*-distance in the NS5B region (0.24) fails to meet the criterion (amino acid *p*-distance > 0.3 for NS5B to the closest relative) (Fig. 2). Consequently, RfHV and HaHV would be classified as the same species under the ICTV standards. However, their hosts are taxonomically substantially different: the host of RfHV, the rusty-margined flycatcher, belongs to the order Passeriformes, whereas the host of HaHV, the Hispaniolan amazon, is in the order Psittaciformes. Additionally, the genome nucleotide sequence identity between RfHV and HaHV is low at 43.2%. Furthermore, exceptions exist within the current ICTV classification of the genus *Hepacivirus. Hepacivirus peromysci* and *Hepacivirus myodae* are classified as separate species despite their *p*-distance of NS5B region being 0.27 (below the ICTV criterion). This classification takes into account that their respective hosts belong to different genera: the host of members of *Hepacivirus peromysci* is the bank vole (*Myodes glareolus*), while that of *Hepacivirus myodae* is the eastern deer mouse (*Peromyscus maniculatus*) [1]. Considering that the bank vole and the eastern deer mouse belong to the same family (Cricetidae), it would be appropriate to classify RfHV and HaHV, whose hosts differ at the order level, as distinct species.

Our results suggest that primarily hosts of RpHV is rock pigeons and RpHV is widely distributed at least within China. Although we screened RNA-seq datasets from diverse bird species, RpHV was detected exclusively in datasets derived from rock pigeons. This suggests that the rock pigeon is the primary host of RpHV. The some RpHV-positive data are from Sichuan and Beijing suggests that RpHV is widely distributed throughout China (Tables 1 and S5). Additionally, although the screened datasets included pigeon samples from China, India, and the UK, RpHV was detected only in samples from China. However, this might reflect sampling bias rather than true geographic restriction. Further molecular epidemiological studies are required to understand the host range and distribution of RpHV.

The RfHV and HaHV lineages may be distributed around the Caribbean region to South America. The HaHV-positive individual was kept in Puerto Rico (Table 1), and Hispaniolan amazon is endemic to the Caribbean island of Hispaniola [40]. Meanwhile, the habitat of Rusty-margined flycatcher is Central America to the South America [41], and the RfHV-positive individual was born in Brazil (BioSample SAMN29332761). These data suggest that RfHV, HaHV, and possibly related viruses may have a natural distribution primarily from the Caribbean region to South America. However, since only two individuals were found positive for this lineage of viruses, further surveillance is required to test this hypothesis.

While all avian hepaciviruses formed a monophyletic group indicating strong host specificity to birds, incongruence between the viral and host phylogenetic trees suggests past cross-species transmission of avian hepaciviruses within birds. Most hepaciviruses identified to date form distinct phylogenetic clades that correspond to the taxonomic class of their hosts (Fig. 3). The hepaciviruses identified in this study also belong to a clade with known avian hepaciviruses, suggesting that this lineage of viruses possesses strong host specificity for birds. Furthermore, some incongruencies were observed between the phylogenetic trees of viruses and their hosts (Fig. 4). This suggests that avian hepaciviruses have caused cross-species transmission in the past. Due to the lack of virological data on avian hepaciviruses, it remains unclear whether the modern hepacivirus can cause cross-species transmission. However, since multiple species of captive birds are often co-housed within the same facility, the risk of cross-species transmission may need to be considered.

This study may have partially elucidated the tissue tropism of the identified hepaciviruses. RpHV was detected in samples derived from subcutaneous fat, visceral fat, and oral mucosa, suggesting the possibility of systemic infection or viremia (Table 1). Meanwhile, RfHV was detected in a liver sample, and HaHV was detected in a blood sample. Thus, viruses of the RfHV and HaHV lineages might exhibit a transmission mode similar to HCV. HCV is hepatotropic and transmitted via blood [42]. Therefore, it is possible that viruses of the RfHV and HaHV lineages are also hepatotropic and transmitted via blood. Further studies are warranted to understand the tropisms and transmission modes of avian hepaciviruses.

The pathogenicity of identified hepaciviruses remains unclear. Among the RpHV-positive datasets, two were derived from diseased individuals (Table S5). However, these two individuals were described to have been used in an infection experiment with *Trichomonas gallinae*, implying that their symptoms may be derived from *T. gallinae*. Furthermore, no data regarding the health status of the RfHV- and HaHV-positive individuals were available. Thus, the pathogenicity of these hepaciviruses is unclear. Further epidemiological, pathological, and ideally experimental infection studies are needed to elucidate the pathogenicity of avian hepaciviruses.

In this study, we identified three novel hepaciviruses from birds. Our findings contribute to a deeper understanding of the diversity of hepaciviruses. However, many aspects of their virological characteristics and pathogenicity remain unclear, highlighting the need for further studies.

## Acknowledgements

The super-computing resources were provided by Human Genome Center, the Institute of Medical Science, the University of Tokyo.

This study was supported by KAKENHI grant numbers JP21H01199 (MH), JP22K19234 (MH), JP23K20902 (MH), JP24K21922(MH), JP25K10374 (TT), JP25K02498 (KM), and JP24K18455 (MK) and by the 2024/2025 Osaka Metropolitan University (OMU) Strategic Research Promotion Project (Young Researcher) (MK), by the Japan Science and Technology Agency (JST) Moonshot R&D under grant number JPMJMS2025 (KM), and by the Japan Agency for Medical Research and Development (AMED) under grant number 25fk021016 (KM).

