## Supplementary Figure for "Identification of novel hepaciviruses in rock pigeon (*Columba livia* [Gmelin, 1789]), rusty-margined flycatcher (*Myiozetetes cayanensis* [Linnaeus, 1766]), and Hispaniolan amazon (*Amazona ventralis* [Statius Muller, 1776])"

**A**

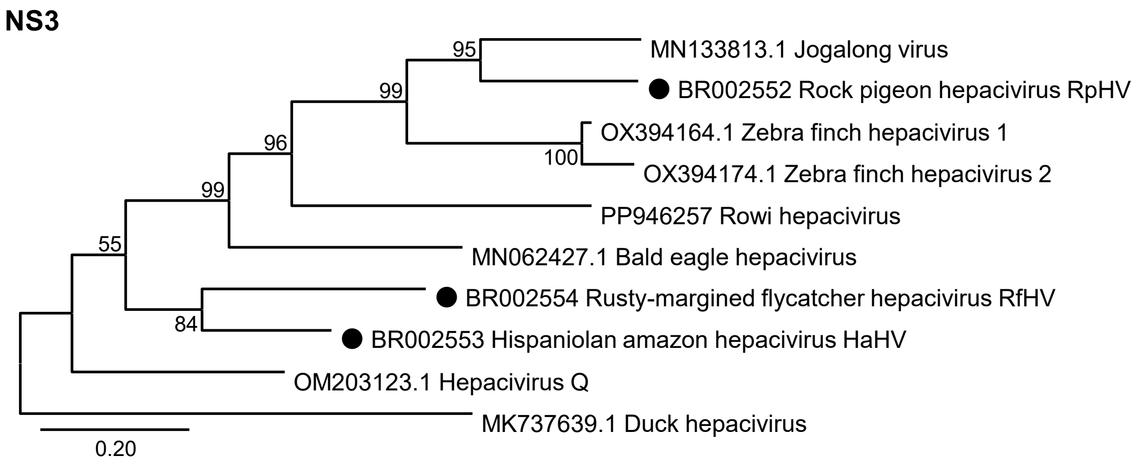


**B**

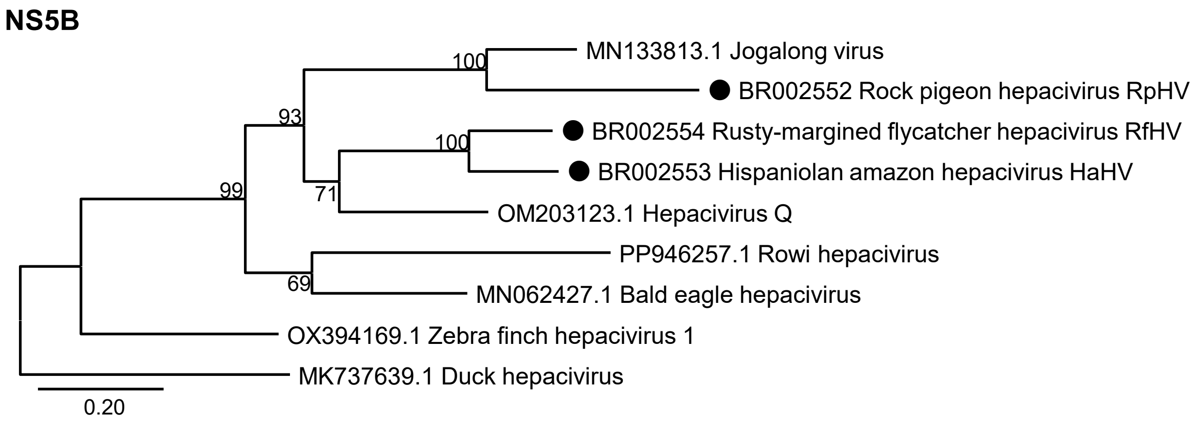


**Supplementary Figure 1. Phylogenetic inference using partial sequences.** Phylogenetic trees were inferred by the maximum likelihood method using amino acid sequences of the NS3 **(A)** and NS5B **(B)** regions of the three novel hepaciviruses identified in this study and known avian and reptile hepaciviruses. The scale bar indicates the number of amino acid substitutions per site. Black circles indicate the novel hepaciviruses identified in this study.
